# A Hypothesis to Explain Cancers in Confined Colonies of Naked Mole Rats

**DOI:** 10.1101/079012

**Authors:** Michael E. Hochberg, Robert J. Noble, Stanton Braude

## Abstract

Naked mole rats (NMRs) are subterranean eusocial mammals, known for their virtual absence of aging in their first 20 to 30 years of life, and their apparent resistance to cancer development. As such, this species has become an important biological model for investigating the physiological and molecular mechanisms behind cancer resistance. Two recent studies have discovered middle and late-aged worker (that is, non-breeding) NMRs in captive populations exhibiting neoplasms, consistent with cancer development, challenging the claim that NMRs are cancer resistant. These cases are possibly artefacts of inbreeding or certain rearing conditions in captivity, but they are also consistent with evolutionary theory.

We present field data showing that worker NMRs live on average for 1 to 2 years. This, together with considerable knowledge about the biology of this species, provides the basis for an evolutionary explanation for why debilitating cancers in NMRs should be rare in captive populations and absent in the wild. Whereas workers are important for maintaining tunnels, colony defence, brood care, and foraging, they are highly vulnerable to predation. However, surviving workers either replace dead breeders, or assume other less active functions whilst preparing for possible dispersal. These countervailing forces (selection resulting in aging due to early-life investments in worker function, and selection for breeder longevity) along with the fact that all breeders derive from the worker morph, can explain the low levels of cancer observed by these recent studies in captive colonies. Because workers in the field typically never reach ages where cancer becomes a risk to performance or mortality, those rare observations of neoplastic growth should be confined to the artificial environments where workers survive to ages rarely if ever occurring in the wild. Thus, we predict that the worker phenotype fortuitously benefits from anti-aging and cancer protection in captive populations.

## Introduction

Despite cancer and cancer-like phenomena being present across the tree of life (Aktipis et al. 2015), incidence data are mostly limited to mammals. Peto was one of the first to describe the surprising observation that body mass and life span show little or no correlation with cancer incidence across species (Peto et al. 1975; Peto 1977). The so-called “Peto’s paradox” (Nunney 1999; Caulin and Maley 2011) is rooted in the expectation that cancer incidence should correlate with cell number (targets for transformation) and organism life span (exposure time per target and/or target turnover). Higher than expected incidence suggests relaxed or compromised tumour prevention, whereas lower than expected levels indicate reinforced prevention and/or suppression.

Natural selection for cancer resistance is a plausible explanation for Peto’s paradox (Caulin and Maley 2011). According to this theory, selection to increase body size and/or longevity will incur costs of increasing vulnerability to invasive neoplasms and decreased fitness due to reduced survival and reproduction. Thus, cancer risk is expected to constrain the evolution of certain life history traits (Kokko and Hochberg 2015). Theory predicts that an increase in life span should have a larger effect on cancer risk than an equivalent increase in body size (e.g. (Nunney 1999)). For example, the simplest multi-stage model predicts that lifetime cancer risk is proportional to *s*(*ud*)^*M*^. Here, *s* is the number of stem cells, *u* is the mutation probability at any one (of a fixed number) of *M* genes that must be mutated for cancer to occur, and *d* is the lifetime number of stem cell divisions. It follows that if three genetic transformations are required, then increasing life span by a factor of 10 changes cancer risk as much as increasing body size by a factor of 1,000. The latter figure remains relatively large (approximately 75) even if we account for the observation that larger animal species generally live longer (assuming that when *s* increases, *d* will increase proportional to *s*^0.2^ (Speakman 2005)).

## Naked mole rats can get cancer

Naked mole rats (*Heterocephalus glaber*) have been proposed as a model species for understanding cancer prevention and suppression. NMR colonies are composed of tens to hundreds of individuals, among which there is usually only one breeding female and one to three breeding males (e.g., (Jarvis et al. 1994)). These organisms have exceptional life spans in captivity relative to their body mass (Sherman and Jarvis 2002; Speakman 2005; Buffenstein 2008), and until recently approximately 2,000 reported necropsies had never revealed an invasive neoplasm (Buffenstein 2008; Grimes et al. 2012). Research to explain this surprising observation and the resistance of NMRs to DNA damage has identified several putative cancer resistance mechanisms (e.g., (Seluanov et al. 2009; Liang et al. 2010; Kim et al. 2011; Tian et al. 2013; Keane et al. 2014; Miyawaki et al. 2016)).

Recently, Delaney and coworkers (Delaney et al. 2016) described the first known cases of cancer in NMRs. The two specimens were male workers (that is, non-breeders) with estimated ages of 20 and 22 years. These specimens came from different colonies, reducing the likelihood of a colony-specific effect such as rearing conditions. These observations are important in what was thought to be a species immune to invasive carcinoma. Although the report does not indicate whether the health of one of the NMR individuals (a 22 year old male) was threatened by the disease, the second specimen (an estimated 20 year old male) was humanely euthanized and presented a series of disorders. In particular, “densely cellular neoplasm was detected, expanding and effacing the gastric submucosa and mucosa” (Delaney et al. 2016). It is therefore likely that had this individual not been sacrificed, cancer would have contributed as a cause of death.

Also recently, Taylor and colleagues (Taylor et al. 2016) presented pathology results from a sample of 72 necropsies from a single NMR colony. They found four confirmed cases of neoplasia and one presumptive case (all workers, Nicolas Milone, pers. Comm). Interestingly, each of the pathologies was located in a different anatomical site (and indeed two individuals showed metastasis), and the mole-rat ages were as young as *c.* 6 years old (more conservative estimate of 11 years old (Taylor et al. 2016)). Regarding the former observation, the authors hypothesize that more than one mutational origin may have been responsible for the range of cancer sites and pathologies, and that a previously described mechanism of cancer resistance (Tian et al. 2013) might not be effective in preventing the types of cancer observed. The latter result indicates that cancer can become apparent in middle-aged individuals and challenges previous work indicating that obvious signs of senescence do not emerge before *c*. 24 year of age (Buffenstein 2008). However, Taylor and coworkers (Taylor et al. 2016) note that rearing conditions, including inbreeding, which occurs only rarely in the wild (Braude 2000; O’Riain and Braude 2001; Braude 2007; Ingram et al. 2015)^1^, may have contributed to the extraordinarily high frequency of cancers in the population studied. Taylor’s study colony is descended from a breeding pair that, in turn, was descended from a colony collected in the field in 1991. Hence, high levels of pedigree inbreeding are to be expected.

The exact causes of the cancers observed in these two studies are unknown. Although certain physiological indices may decrease moderately in NMR workers after 5 years of age (O’Connor et al. 2002), signs of reduced physiological performance and lower reproductive success, should they occur, happen only after about 20 to 24 years of age, depending on which index is measured (summarized in (Buffenstein 2008; Edrey et al. 2011)). Cohort data for captive colonies show that mortality rates are low and constant until NMRs are in their mid-twenties, which has been interpreted as evidence for negligible aging, with a sudden onset of senescence at the end of life (Buffenstein 2008). This pattern of mortality results in a “type I” survivorship curve (Begon et al. 2005), with a life expectancy at birth of a captive worker NMRs of approximately 20 years (Figure 3 in (Buffenstein 2008)). Data from other laboratory colonies suggest negligible differences in maximum life spans between workers and breeders (Buffenstein 2008). Finally, molecular data support mortality observations that individuals exhibit negligible senescence (Perez et al. 2009; Kim et al. 2011). Unfortunately detailed life-table analyses of physiological indices, performance measures, and survival and reproduction, comparing workers and breeders are yet to be conducted.

## Evolutionary theory of aging and its application to NMRs

Based on what is known about NMR biology and ecology we argue that evolutionary theories of aging are consistent with the finding that naked mole rats have specific adaptations that prevent morbidity and mortality from cancer (Seluanov et al. 2009), but that occasional cancers may nevertheless occur in captive middle-aged and older workers, as reported by both Taylor et al and Delaney et al.

Evolutionary theory predicts a progressive decrease in the strength of selection with age, which in turn can explain aging and senescence (Medawar 1952; Williams 1957; Hamilton 1966). In eusocial species however, sterile castes have significantly shorter lifespans compared to reproducing individuals (Alexander et al. 1991; Keller and Genoud 1997; Chapuisat and Keller 2002; Heinze and Schrempf 2008; Kramer and Schaible 2013). This can be explained if sterile workers in eusocial species express performance-enhancing genes early in life (Finch 1990; Keller and Jemielity 2006), leading to high-risk, high-payoff behaviours that increase the inclusive fitness of the genes associated with these behaviours (Queller 1996; Bourke 2007), but that may entail costs to lifespan compared to reproducing individuals (i.e., antagonistic pleiotropy (Williams 1957)). Analogously, the disposable soma theory suggests that, due to a limited energy budget, investments in somatic maintenance (e.g., sterile worker longevity) are only selected insofar as reproductive output of the higher individual increases (Kirkwood 1977; Kirkwood and Austad 2000). Although a comprehensive theory of ageing is yet to be developed for eusocial species, an expectation based on senescence theory in age-structured populations (e.g., (Hamilton 1966; Abrams 1993; Williams and Day 2003; Caswell and Shyu 2016)) is that increased early extrinsic mortality and decreased late mortality should select for reduced senescence (Caswell 2007), but it is not clear in the context of eusocial species how this may affect the absolute and relative lifespans of sterile and reproducing castes.

Although data on NMR survivorship is limited, it is consistent with selection against early pleiotropic deleterious alleles (e.g., aging, cancer). When predation, parasites, pathogens and failed dispersal are eliminated as sources of mortality for captive populations, both workers and breeders exhibit extraordinary lifespans (Sherman and Jarvis 2002). Data from captive NMR colonies suggest that workers perish at a more or less constant annual rate of *c*. 10%–15% per year up to 24 years of age (Figure 4 in (Buffenstein 2008)). Although the annual mortality rates of breeders prior to 20 years of age are unknown, limited data from (a possibly inbred) field-collected individuals that were reared in captivity suggest that workers and breeders differ little in their survival rates until old age (Sherman and Jarvis 2002).

But does this picture of equally low mortality in breeders and workers in captivity accurately portray how selection operates in the wild? The answer appears to be no. In the wild, breeders have been documented at up to 17 years of age, but most workers disappear from their natal colonies by 2-3 years of age, either from disease or predation (see below). Those workers reaching approximately 2 to 3 years of age become dispersers (O’Riain and Braude 2001) and either reside in the colony in a less active state, building fat reserves, or leave the natal colony to potentially establish new colonies or join existing ones (O’Riain et al. 1996). Jarvis (1991) referred to these large, older animals as “non-workers” (although she included breeding males under this umbrella term). High rates of predation mortality are suffered by dispersers during failed attempts to emigrate and establish new colonies (Braude 2000; O’Riain and Braude 2001; Braude et al. 2001). By capturing entire colonies of naked mole-rats, marking individuals and re-trapping many colonies year after year from 1985 through 2004 (Braude and Ciszek 1998), we have found that presence in the colony approximates a “type II” survivorship curve for workers and dispersers in the wild (Figure 1; Table 1), rather than the type I curve that is typical of captive mammals (and of modern humans). The annual loss rates from colonies (i.e., mortality within the colony plus dispersal from the colony) are 60% (58-62% with standard errors) for males and 56% (55-58%) for females. Therefore, residence expectancies in natal colonies are 1/0.60 = 1.7 (1.6-1.7) years and 1/0.56 = 1.8 (1.7-1.8) years, respectively (O’Riain and Braude 2001). Although the fraction of these animals that perish within the colony or in the actual process of dispersing is unquantified, very few dispersers got to the point of establishing even a little colony. From field data between 1987 and 2000 there were 25 such nascent colonies (out of 8000 NMRs captured), with three or fewer animals, and none of these colonies persisted for more than a year or grew to a size of more than 12 individuals (S. Braude, *pers. obs.*).

**Figure 1.**
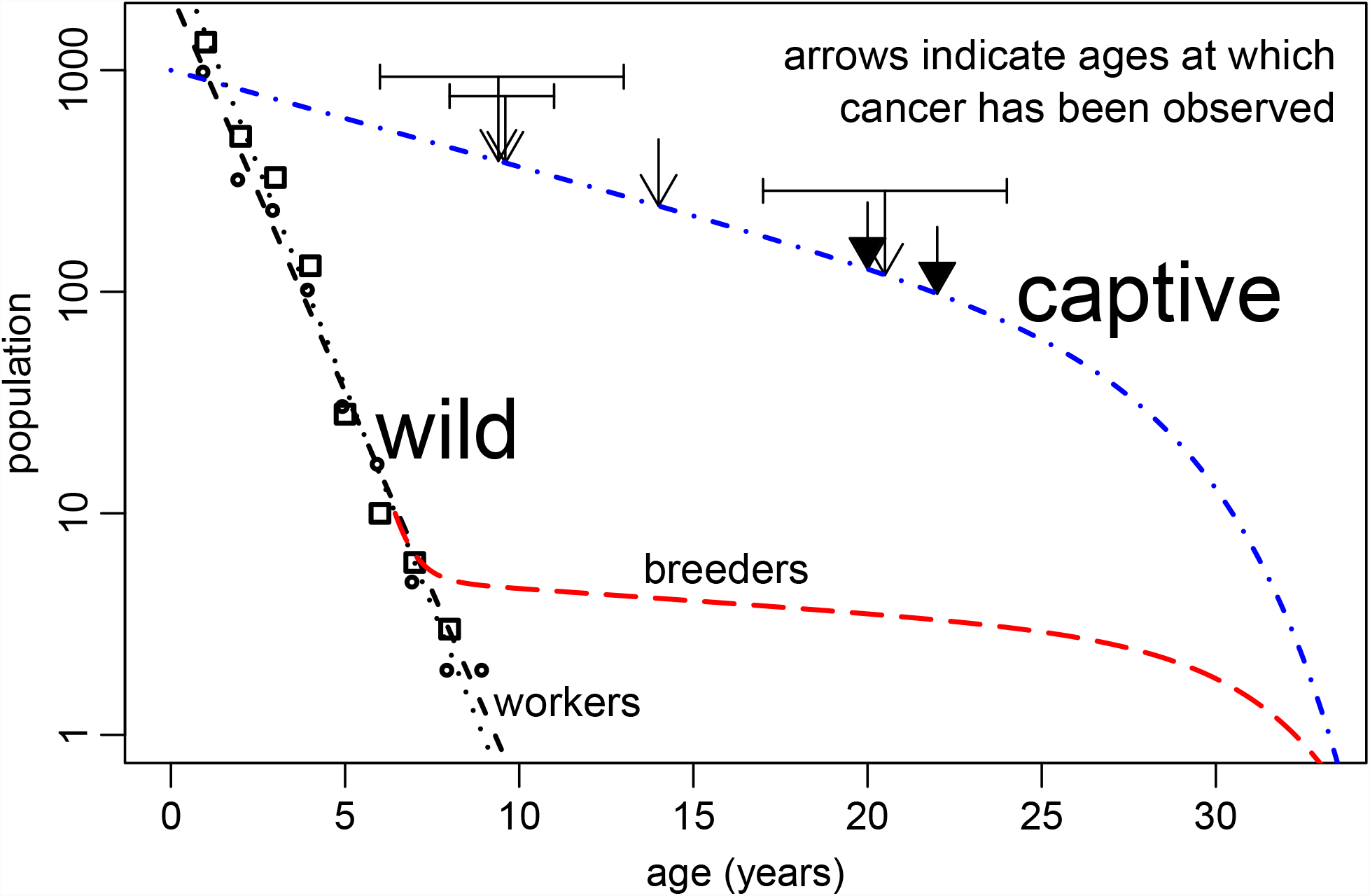
Residence curves for naked mole rat colonies in the wild and survival curve for captive naked mole rats. Regression analysis of field data (Table 1) shows that presence in wild colonies decreases approximately logarithmically for both males (squares and black dotted line) and females (circles and black dashed line). This "type II” survivorship pattern indicates that most mortality and dispersal is independent of age (although unquantified, most individuals are likely to have perished (O’Riain and Braude 2001)). Contrastingly, in captive populations, mortality rates are relatively low until approximately 25 years of age and then increase sharply (e.g., (Buffenstein 2005; Buffenstein 2008)) (blue dashed and dotted line), corresponding to a “type I” survivorship curve. The same type of survivorship curve has been observed in captive breeders and is hypothesised to apply to breeders in the wild (red dashed line), which include a single queen, and typically 1 to 3 breeding males. Arrows indicate the ages at which cancer has been observed in captive naked mole rat workers by Delaney et al. (solid arrowheads) and Taylor et al. (open arrowheads), with error bars indicating minimum and maximum estimates.

**Table 1:**
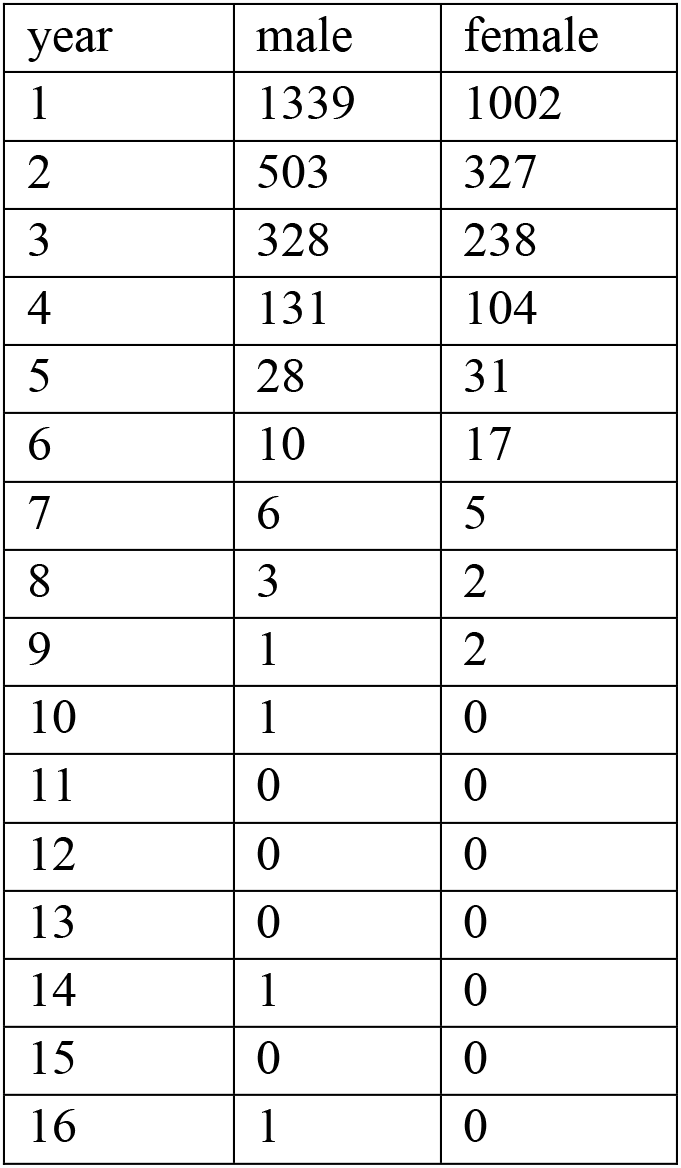
Residence of naked mole rats in wild colonies over time. Seventy-nine entire colonies in Neru National Park, Kenya were captured multiple times between 1986 and 2004. Each individual was measured, marked, and returned to its burrow. The table includes only individuals judged to have been less than 1 year old at the time of first capture, either because they weighed less than 23 grams or because they had not been caught during a complete colony capture in the preceding year. Population decreases are due to death and emigration. The expected survival time for an emigrant is relatively very short.

## A hypothesis to explain the observations

We propose that the unusually low rate of cancer in naked mole rats results from the balance of different selective forces on risky functions (worker morphs and disperser morphs) and on protected functions (breeders), along with the developmental necessity that breeders are always derived from either workers or dispersers.

The eusocial system in NMRs and their ecology, together with the above laboratory and field observations suggest how contrasting forces of natural selection have resulted in reinforced, but possibly not perfect, cancer protection *in captive colonies* (Figure 2). NMRs exhibit age-polyethism, whereby different colony functions and to some extent associated morphological features change with age (Lacey and Sherman 1991). Foraging, tunnel maintenance and colony protection are crucial in this species in the wild, and are conducted to a great extent by young workers (0-2 years of age). Worker traits that contribute to colony fitness (colony survival and the establishment of new colonies) should be under selection (Queller 1996); selection for risky, early performance in eusocial species can result in late-expressed pleiotropic costs in worker phenotypes (Oster and Wilson 1978). However, NMR eusociality differs importantly from the better-studied cases in many eusocial insects, in that workers are not dead-end steriles as, for example, in certain ants and in honey bees: reproductive NMRs are replaced by physiologically mature workers in NMRs (Jarvis 1991), rather than by brood (e.g., larvae in most social insects). This means that selection on certain worker traits is more linked to selection on breeder traits than in eusocial species with dead-end steriles. As such, the two main traits under selection are: (1) traits specific to workers that influence inclusive fitness (protecting the colony, maintaining tunnels, brood care), and (2) traits shared by workers and breeders that more directly influence reproductive output, namely survival to reach breeding status (Lacey and Sherman 1991; Lacey and Sherman 1997) and survival and reproduction once becoming a breeder (crucial for social stability (Jarvis 1991; Clarke and Faulkes 1997)). The latter feature is favoured by additional selection against senescence associated with pronounced mortality in the first months/years of life (e.g., (Caswell and Shyu 2016) and data from the present study), the relatively highly protected environment of breeders in lower subterranean burrows (Hamilton 1978; Alexander et al. 1991; Keller and Genoud 1997; Keller 1998), and by age-dependent increases in reproductive output (Stearns 1992; Buffenstein 2008). Thus, early worker performance, if contributing sufficiently to colony fitness, could be associated with some senescence, but this will be limited due to selection for longevity in these same individuals that reach breeding status.

**Figure 2.**
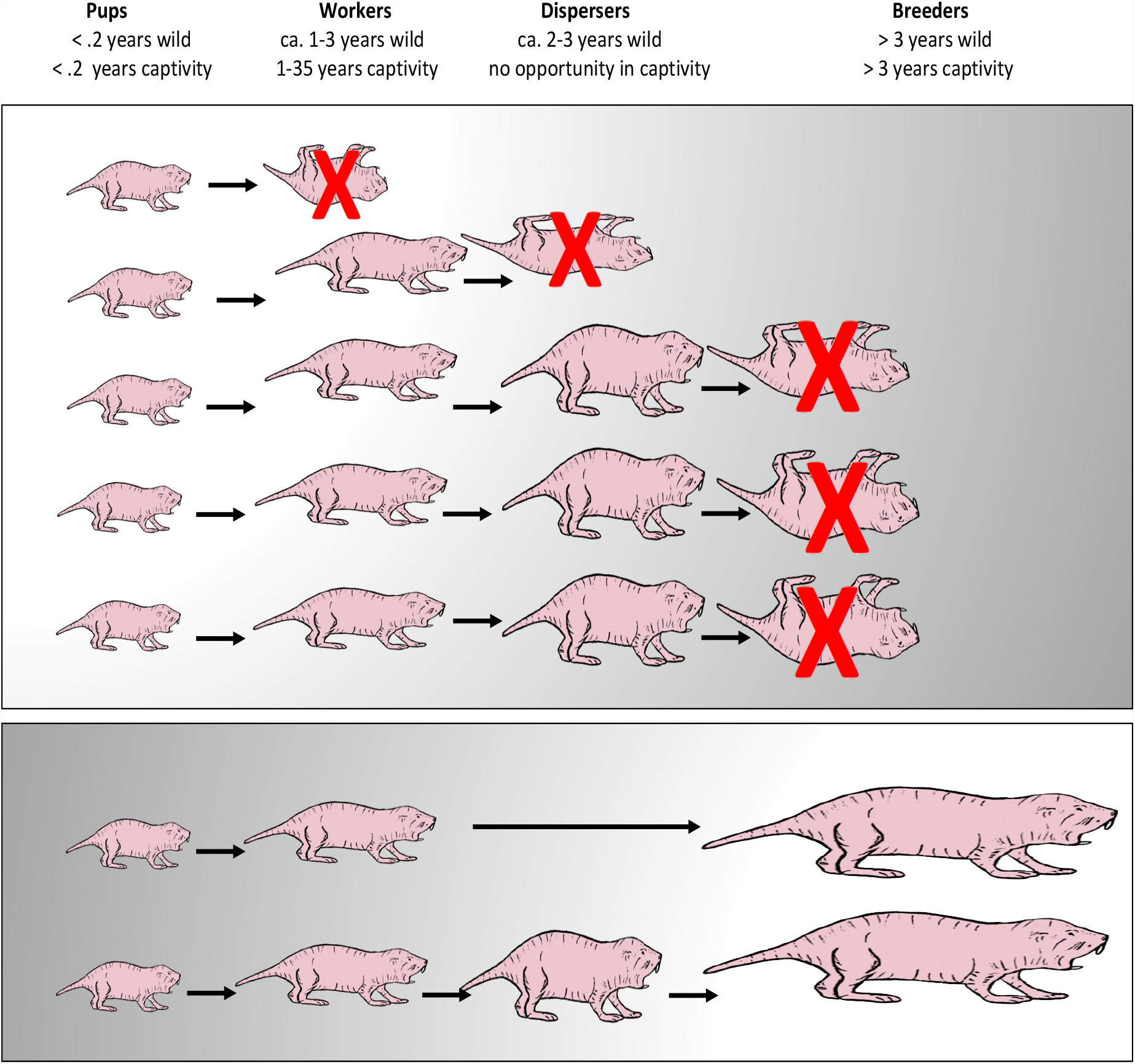
Divergent life trajectories of wild and captive naked mole rats. **Top Panel:** Early mortality of pups, workers, and especially dispersers, in the wild should select against pleiotropic genes that would lead to earlier senescence and cancer. This does not completely occur, however, because of selection for early performance (with later, pleiotropic effects). The countervailing forces of early mortality and selection for early performance result in a low degree of aging and occasional cancers that affect performance. **Bottom Panel:** However, all pups, workers and dispersers retain the biological potential to mature, and have a finite probability of surviving and becoming a reproductive. The safety of queens and breeding males in the lower burrows selects strongly for longevity and cancer resistance genes. See main text for further explanation.

Alexander and coworkers (Alexander et al. 1991) argued that relieving a reproducing social insect from high-risk tasks would select for longer life spans in the reproductive phenotype. It is therefore tempting to speculate that worker NMRs should show greater aging (and cancer) than breeders, because the former perform most of the risky functions in the colony. However, as noted above and unlike social insects where differences in aging between castes has been found (Chapuisat and Keller 2002; Keller and Jemielity 2006; Lucas et al. 2016), NMR breeders derive from workers and dispersers, meaning that much of the adaptive gene expression in the former is likely to be similar in the latter two morphs. This is effectively what the data show (negligible cancer incidence) when workers are reared beyond ages attained in the wild.

Despite a similar ecology and the independent evolution of anti-aging and anti-cancer mechanisms (Nasser et al. 2009; Manov et al. 2013), it is interesting that cancers have never been observed in the blind mole rat (*Spalax sp*.; BMR), a solitary species that is phylogenetically closer to mice than to NMRs. Assuming that the observations of cancers in captive NMRs by Delaney et al and Taylor et al are not artefacts of inbreeding or particular rearing conditions and that the risk of cancer in BMRs is indeed lower than that in NMRs, then this lends support to our hypothesis. Namely, the eusocial system in NMRs results in a low level of aging and cancer in workers as an *artefact* of workers surviving in captive colonies beyond ages possible in the wild.

## Conclusion and future directions

Evolutionary theory can explain the observation of rare cancers in captive naked mole rat workers. Nevertheless, these observations of cancer are, in effect, artefacts of ecological disruption (Hochberg and Noble, submitted), insofar as they occur in animals that would never attain the same ages in the wild.

Due caution is necessary in interpreting our predictions, since the majority of observations in captivity involve worker NMRs only, and factors contributing to cancer such as rearing conditions and inbreeding are unknown (for other effects of inbreeding in this species, see e.g., (Ross-Gillespie et al. 2007)). Future work should develop and analyse explicit mathematical models of aging and cancer for a range of solitary and eusocial biologies that take into account genetics, demography, inbreeding, and differential gene expression. Equally important, experimental study is needed to compare and contrast survivorship and aging in other long-lived species belonging to the same clade as NMRs, that is, the suborder Hystricomorpha (e.g., other African mole rats, chinchillas, and old world porcupines). Finally, research should compare the genetics and gene expression of wild caught NMRs and laboratory colonies, to assess the possible contribution of pedigree inbreeding to observed cancers in laboratory colonies.

## Acknowledgements

We are indebted to Ophélie Ronce, Hanna Kokko, Laurent Keller, Daniel Promislow and David Queller for constructive comments on the manuscript and to Chris Faulkes, Kyle Taylor and Nicolas Milone for discussions. MEH acknowledges support from the Agence National de la Recherche (EvoCan ANR-13-BSV7-0003-01), the CNRS (APICS 06313), and the Institut National du Cancer (2014-1-PL-BIO-12-IGR-1). Field work in Kenya (Meru National Park) was approved by the Office of the President of Kenya (field permit number 15C/116), and by Washington University IACUC (Animal ethics number A-3381-01).

## Footnotes

1 NMR throughout almost all of their range are not inbred. However, many of the captive colonies were bred from animals at the southern frontier of the distribution where a bottleneck event occurred 100 years earlier (Ingram et al. 2015). In addition, captive breeding programs for this species accidentally promote inbreeding as the system of mating.

